# Rubisco supplies pyruvate for the 2-*C*-methyl-D-erythritol-4-phosphate pathway

**DOI:** 10.1101/2024.01.28.577108

**Authors:** Sonia E. Evans, Yuan Xu, Matthew E. Bergman, Scott A. Ford, Yingxia Liu, Thomas D. Sharkey, Michael A. Phillips

**Author notes:** Current address: Department of Biochemistry, Purdue University, West Lafayette, IN, 47907-2063, USA.

## Abstract

Ribulose-1,5-bisphosphate carboxylase/oxygenase (rubisco) produces pyruvate in the chloroplast through beta-elimination of the aci-carbanion intermediate. Here we show that this side reaction supplies pyruvate for isoprenoid, fatty acid, and branched chain amino acid biosynthesis in photosynthetically active tissue. 13C labeling studies of whole Arabidopsis plants demonstrate that the total carbon commitment to pyruvate is too large for phosphoenolpyruvate (PEP) to serve as precursor. Low oxygen stimulates rubisco carboxylase activity and increased pyruvate production and flux through the 2-C-methyl-D-erythritol-4-phosphate (MEP) pathway, which supplies the precursors for plastidic isoprenoid biosynthesis. Metabolome analysis of mutants defective in PEP or pyruvate import further supported rubisco as the main source of pyruvate in chloroplasts. Rubisco beta-elimination leading to pyruvate constituted 0.7% of the product profile in in vitro assays, which translates to 2% of the total carbon leaving the Calvin-Benson-Bassham cycle (CBC). These insights solve the so-called pyruvate paradox, improve the fit of metabolic models for central metabolism, and connect the MEP pathway directly to carbon assimilation.

## Main text

Ribulose-1,5-bisphosphate carboxylase/oxygenase (rubisco) produces pyruvate in the chloroplast through β-elimination of the *aci*-carbanion intermediate^1^. Here we show that this side reaction supplies pyruvate for isoprenoid, fatty acid, and branched chain amino acid biosynthesis in photosynthetically active tissue. ^13^C labeling studies of whole Arabidopsis plants demonstrate that the total carbon commitment to pyruvate is too large for phosphoenolpyruvate (PEP) to serve as precursor. Low oxygen stimulates rubisco carboxylase activity and increased pyruvate production and flux through the 2-*C*-methyl-D-erythritol-4-phosphate (MEP) pathway, which supplies the precursors for plastidic isoprenoid biosynthesis^2,3^. Metabolome analysis of mutants defective in PEP or pyruvate import further supported rubisco as the main source of pyruvate in chloroplasts. Rubisco β-elimination leading to pyruvate constituted 0.7% of the product profile in *in vitro* assays, which translates to 2% of the total carbon leaving the Calvin-Benson-Bassham cycle (CBC). These insights solve the so-called ‘pyruvate paradox’^4^, improve the fit of metabolic models for central metabolism, and connect the MEP pathway directly to carbon assimilation.

The MEP pathway supplies the precursors for plastid-derived isoprenoids, a large, diverse group of products involved in photosynthesis^5^, defense^6^, and communication^7^. In the committing step of the MEP pathway in plants, 1-deoxy-D-xylulose-5-phosphate synthase (DXS) condenses pyruvate and glyceraldehyde-3-phosphate (GAP) into 1-deoxy-D-xylulose-5-phosphate (DXP), followed by conversion to isopentenyl and dimethylallyl diphosphate (IDP and DMADP) in six subsequent steps. IDP and DMADP formed collectively through the plastidic MEP pathway and cytosolic mevalonate pathway are the universal precursors to isoprenoid pigments^8^, electron carriers^9^, phytosterols^10^, and secondary metabolites^11^.

Flux through the MEP pathway depends on GAP and pyruvate availability for the DXS reaction. While the CBC supplies GAP in photosynthetic tissue, the origin of pyruvate has been attributed to alternative sources^12-15^. Compared to other central metabolites, total cellular pyruvate labels slowly in isotopic labeling experiments^16,17^, and large pools may be sequestered in other plant cell compartments that are metabolically inactive during the day. In the cytosol, glycolysis is the source of pyruvate, but in illuminated chloroplasts, two steps of glycolysis (phosphoglycerate mutase and enolase; PGM and ENO) are thought to be downregulated beyond the seedling stage^18^, presumably to prevent catapleurosis of CBC intermediates (Fig 1a). This would effectively block direct formation of pyruvate in photosynthesizing chloroplasts, which is required not only for the MEP pathway but for fatty acid synthesis and the branched chain amino acid pathway as well. The cryptic nature of pyruvate biogenesis, labeling and transport has been described as the ‘pyruvate paradox’^4^.

**Fig 1.**
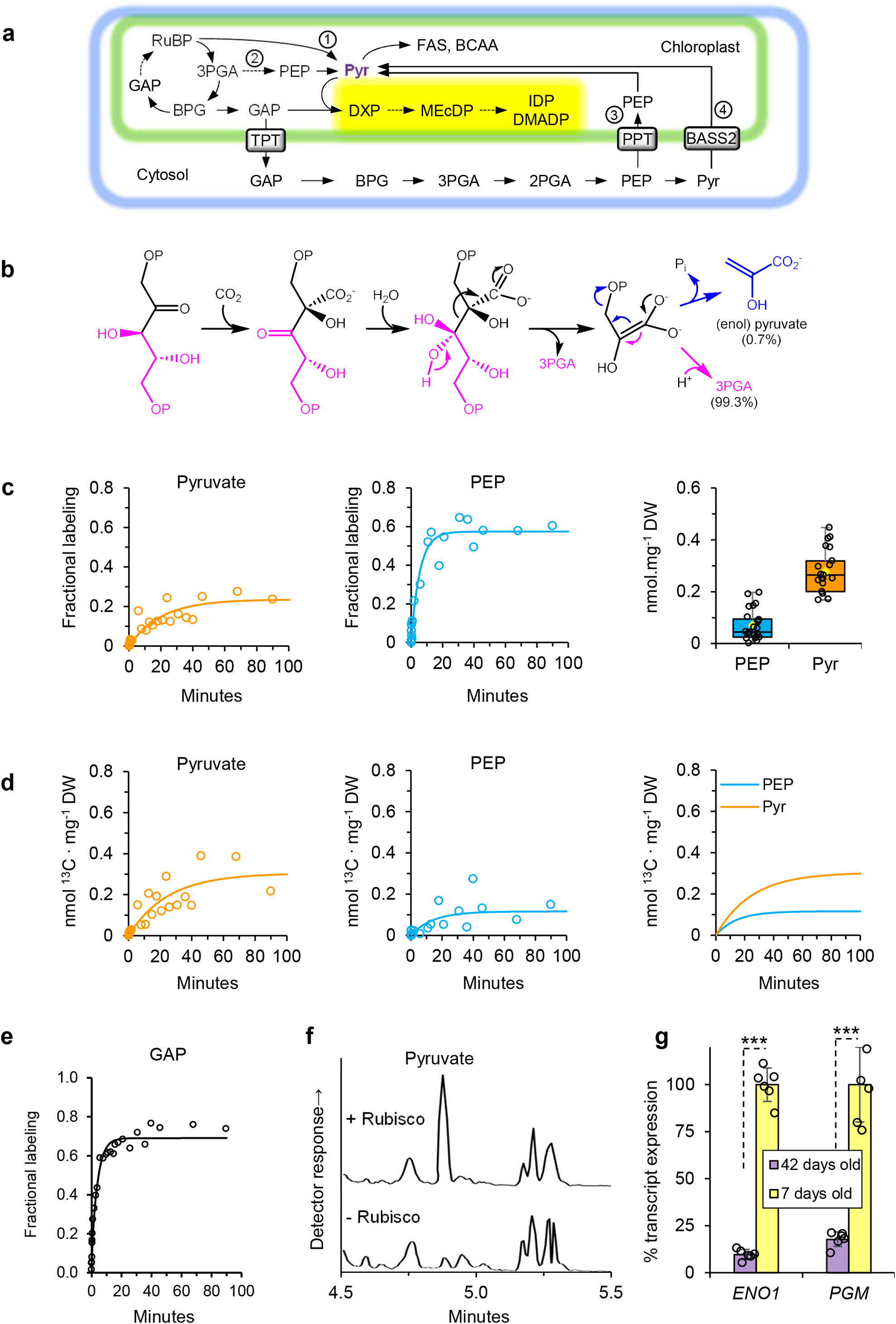
Role of rubisco in supplying pyruvate in illuminated chloroplasts. **a**, Simplified view of central metabolism in leaf tissue emphasizing the potential sources of pyruvate and its role in biosynthetic routes. Abbreviations: RuBP, ribulose-1,5-bisphosphate; 3PGA, 3-phosphoglycerate; 2PGA, 2-phosphoglycerate; PEP, phosphoenolpyruvate; BPG, 1,3-bisphosphoglycerate; GAP, glyceraldehyde-3-phosphate; Pyr, pyruvate; DXP, 1-deoxyxylulose-5-phosphate; MEcDP, 2-C-methylerythritol-2,4-cyclodiphoshate; IDP, isopentenyl diphosphate; DMADP, dimethylallyl diphosphate; FAS, fatty acid synthesis pathway; BCAA, branched chain amino acid pathway; TPT, triose phosphate/phosphate translocator; PPT, phosphoenolpyruvate/ phosphate translocator; BASS2, sodium/pyruvate symporter. The yellow box indicates the 2C-methyl-erythritol 4 phosphate pathway. Dashed line indicates multiple enzymatic steps. **b**, Mechanism of rubisco (as per Andrews and Kane^1^), emphasizing the departure of the ‘lower’ 3-PGA (pink) and the ‘upper’ C_3_ unit, which may depart as 3-PGA (pink) or alternatively as enolpyruvate through β–elimination of the *aci*-carbanion intermediate (blue), which rapidly isomerizes to pyruvate. **c**, Time course label incorporation into the C_3_ central metabolites pyruvate and PEP in whole Arabidopsis rosettes by administering ^13^CO_2_ under physiological conditions. The total cellular pyruvate and PEP pool sizes are shown at right as box plots, with the central line indicating the median. Upper and lower bounds represent quartile 3 (75th percentile) and quartile 1 (25th percentile), respectively, and the whiskers denote the minimal and maximal values of the data points. Mean values (yellow diamond) are representative of n=15-28 individual plants. **d**, Total substrate commitment into pyruvate and PEP pools as determined by: fractional labeling × pool size × #C atoms. An overlay of their fitted exponential curves is shown at right. For curve fitting parameters, see supplementary tables 1 and 2. e (PEP). **e**, Time course label incorporation into glyceraldehyde-3-phosphate in whole Arabidopsis rosettes. **F**, Gas chromatography – mass spectrometry chromatogram showing detection of derivatized pyruvate produced by rubisco in an in vitro enzyme assay with RuBP (see Methods for details). **g**, Transcript of abundance of plastid localized ENOLASE1 (ENO1; At1g74030) and PHOSPHOGLYCERATE MUTASE (PGM; At1g22170) in 7-day old Arabidopsis seedlings and rosette-staged leaves (42 days). Expression of both genes in the seedling is normalized to 100%. APT1 was used as the reference gene. Bar height indicates the mean of n=6 biological replicates analyzed in triplicate (error bars show standard error of measurement). Asterisks indicate significant differences at *** P < 0.0001, as determined by Student’s two-tailed t-test.

In embryos and developing seedlings, reimport of phosphoenolpyruvate (PEP) through the PEP/phosphate transporter (PPT) is considered to be the main source of plastidic pyruvate for anabolic metabolism^19-22^ (Fig. 1a). However, an Arabidopsis mutant of the *PPT1* gene, deemed *chlorophyll a/b binding protein underexpressed1* (*cue1*), displays normal levels of fatty acids in leaves^23^, suggesting that in autotrophic tissue, stromal pyruvate may derive from another source. The *cue1* phenotype was complemented by plastid expression of pyruvate:phosphate dikinase (PPDK), which generates PEP from pyruvate, phosphate, and ATP^24^. This verified that pyruvate is present in the stroma at a concentration sufficient to supply the shikimate pathway with PEP independently of PPT1 when PPDK is overexpressed in this compartment.

Direct pyruvate import has also been invoked to explain its availability in the chloroplast (Fig 1a). A pyruvate/Na^+^ symporter (BILE ACID:SODIUM SYMPORTER FAMILY PROTEIN 2; BASS2) imports pyruvate from the cytosol into the plastid at early developmental stages^25^, but its physiological importance in photosynthetic source tissue remains untested. Andrews and Kane showed that ribulose-1,5-bisphosphate carboxylase/oxygenase (rubisco) produces pyruvate as a minor side product (∼0.7%) under *in vitro* conditions^1^ (Fig 1b). Due to its extremely high rate of carboxylations, even this small fraction could represent sufficient pyruvate to satisfy the cell’s needs for isoprenoid, fatty acid, and branched chain amino acid biosynthesis. Here we sought to compare the relative contributions of PEP or pyruvate import, glycolytic flux, and pyruvate production by rubisco to the stromal pyruvate pool available for the biosynthesis of isoprenoids and other pyruvate-derived central metabolites in the chloroplast.

We examined transcript levels for the plastidic isoforms of PGM and ENO and observed ∼90% reduction in transcripts in adult Arabidopsis rosette tissue compared to 7-day old seedlings (Fig. 1g), consistent with previous reports describing a major downregulation of glycolysis in illuminated chloroplasts^26^. Whole Arabidopsis plants were then subjected to short-term labeling with ^13^CO_2_ under physiological conditions to investigate trafficking of C_3_ intermediates of central metabolism. Mass spectrometry analysis of the pyruvate and PEP extracted from leaf tissue indicated that while PEP was quickly labeled by ^13^C, reaching a plateau of ∼50% enrichment within a few minutes, pyruvate labeled more slowly, reaching a plateau of enrichment of 25% over the course of more than one hour (Fig. 1c). GAP showed the fastest labeling kinetics under these conditions, reaching a plateau of 63% in minutes, and significant label incorporation was evident at the shortest time point our experimental system permitted (∼ 5 s) (Fig. 1e). The comparatively slow kinetics in pyruvate labeling (half-life ∼ 20 min) initially suggested that PEP turnover was sufficient to account for its role as precursor to pyruvate. However, when both fractional labeling plateaus and pool size were taken into consideration (Fig. 1d), the total carbon commitment to pyruvate was ∼3-fold larger than that of PEP (∼0.3 nmol ^13^C atoms · mg^−1^ leaf tissue D.W. for pyruvate vs. ∼0.1 nmol ^13^C atoms · mg^−1^for PEP). This result indicated that total carbon flow into PEP could not account for the larger volume of carbon entering the metabolically active pyruvate pool in the chloroplast. This verified most pyruvate is formed independently of PEP in illuminated chloroplasts.

We tested whether rubisco could supply pyruvate directly in the stroma to account for this pyruvate use by examining the product profile of purified rubisco from *Nicotiana tabacum* incubated with RuBP. Assays were incubated with or without rubisco for 60 min at 25°C (Fig. 1f). Recovery of added RuBP was 88% in the absence of rubisco, while recovery of 3-phosphoglycerate (3PGA) plus pyruvate and residual RuBP was 93% in the presence of rubisco. The difference was not statistically significant. The small amounts of pyruvate and 3-PGA in the minus rubisco treatment was presumed to result from contamination or nonenzymatic conversions. Subtracting these amounts from the + rubisco samples, and considering (PGA + pyruvate)/2 as total carboxylations, an average of 0.71% pyruvates per carboxylation were observed (Supplementary Table 1), with a 95% confidence interval from 0.57 to 0.85%, consistent with Andrews and Kane ^1^. Given that each pyruvate represents three carbons, pyruvate accounted for approximately 2% of the carbon leaving the CBC.

To further test the idea that rubisco carboxylase activity was responsible for pyruvate in the chloroplast, we quantified its capacity to form pyruvate *in planta* under low O_2_ conditions which are known to suppress photorespiration^27^. We hypothesized that an increase in carboxylation rate would proportionally enhance production of pyruvate and DXP. When labeled following acclimation to a low oxygen (1%) atmosphere, wild-type plants displayed the expected decrease in labeling of photorespiratory intermediates, although changes in absolute pool size were minor (Supplementary Fig. 2 and Supplementary Table 2). Low O_2_ conditions provoked an increase in both 3PGA and pyruvate concentration as well as its rate of ^13^C label incorporation (Fig. 2a). The rate of labeling of DXP also increased under low O_2_ conditions (*k* = 0.264 vs. 0.121 for normal O_2_; Fig. 2b and Supplementary table 3), but its labeling plateau and concentration were unchanged. In contrast, suppression of photorespiration increased the labeling plateau and pool size of the MEP pathway intermediate MEcDP and the pathway end products IDP and DMADP without altering their rates of label incorporation (Fig. 2b). The change in the rate of DXP turnover, but not in its pool size, likely reflects the role of DXS as the major rate determining step of the MEP pathway^28^, which increases the rate of flux through the pathway when substrate concentrations rise. DXS is also under allosteric^29,30^ and proteolytic regulation^31^. The subsequent step, catalyzed by DXP reductoisomerase (DXR), is evidently able to keep up with the increased rate of DXP production and maintain DXP levels constant. In contrast, this increase in DXS activity shifts the rate limiting step downstream to 4-hydroxy-3-methylbut-2-enyl diphosphate synthase (HDS), causing an accumulation of its substrate 2-C-methyl-D-erythritol-2,4-cyclodiphosphate (MEcDP), a well-known consequence of increased DXS activity^28,32^. The increase in flux through the MEP pathway when the rubisco carboxylation rate is stimulated by low O_2_ is consistent with a role for rubisco is supplying pyruvate needed for the DXS reaction.

**Fig 2.**
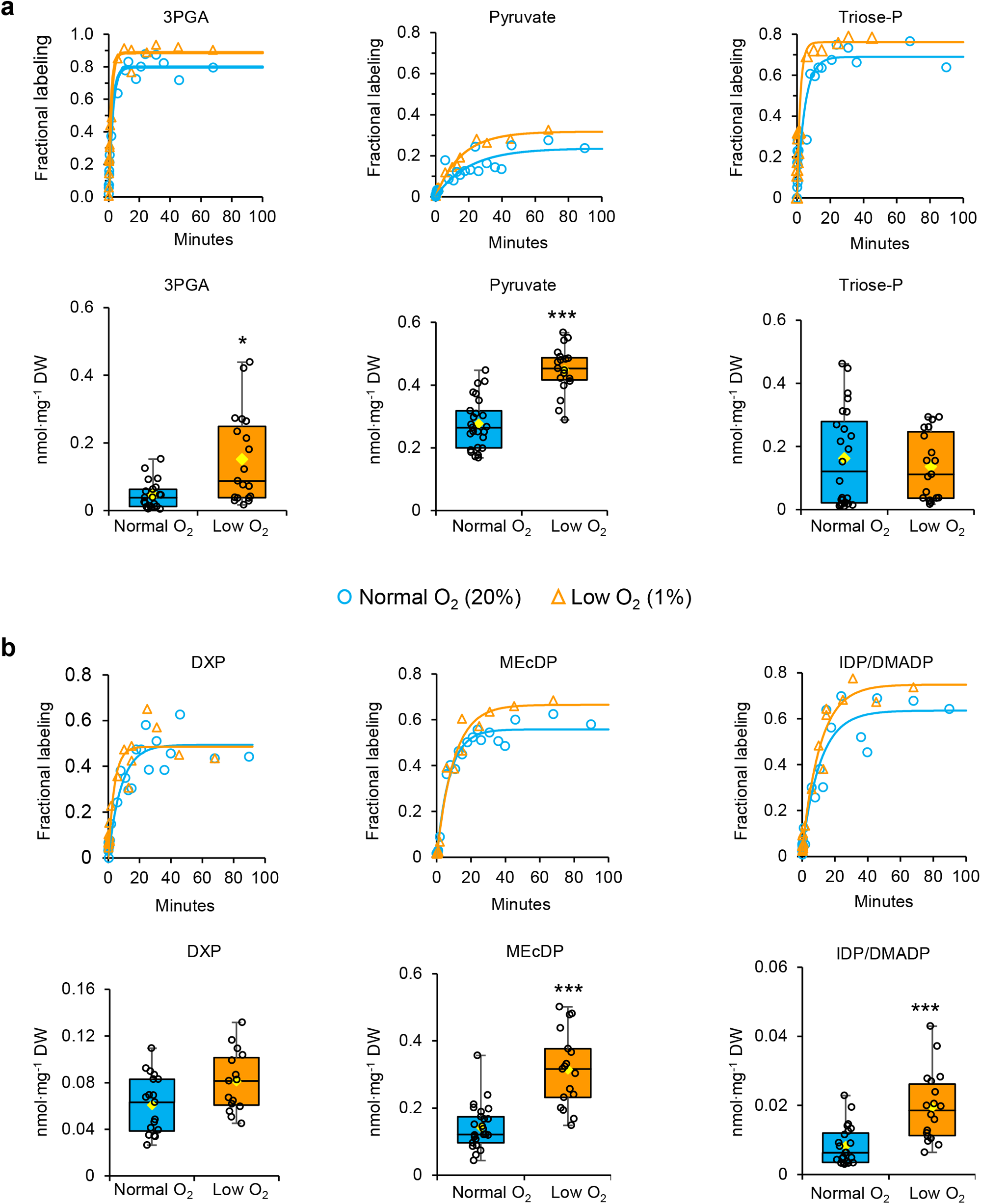
Changes to flux in C_3_ central metabolites and the 2-C-methyl-D-erythritol-4-phosphate (MEP) pathway under low oxygen conditions. **a**, Changes to the ^13^C label incorporation rate and pool sizes of C_3_ intermediates in Arabidopsis leaves under normal (20%) or low (1%) O_2_ conditions. b, Changes to the ^13^C label incorporation rate and pool sizes of MEP pathway intermediates in Arabidopsis leaves under normal (20%) or low (1%) O_2_ conditions. For a and b, each time point represents an individual Arabidopsis plant labeled with 400 μL·L^−1 13^CO_2_ under physiological conditions from 5 s to 90 min (n = 15-28 plants). Each metabolite concentration was determined by comparison to external standard curves. Recovery was corrected by normalization to the internal standard 2-deoxyglucose-6-phosphate. For box plots, the central line separates the median, upper and lower bounds representing quartile 3 (75th percentile) and quartile 1 (25th percentile), respectively. The whiskers denote the minimal and maximal values of the data points. Mean values are displayed as yellow diamonds (n=15-28 plants). Asterisks indicate significant differences at ** P <0.01, *** P < 0.0001, as determined by Student’s two-tailed t-test.

Multiple translocators have been implicated in the provision of pyruvate to the chloroplast, and we examined differences in labeling and pool size in the corresponding Arabidopsis mutants to determine their contribution to this process. The *cue1* mutant cannot import PEP into the chloroplast^22^, and its variegated phenotype (Fig. 3a) is largely the result of an impaired shikimate pathway and deficiency in aromatic amino acid (AAA) biosynthesis^24^. We reasoned that if reimported PEP were the source of pyruvate for isoprenoid biosynthesis, then DXP levels and labeling ought to be highly reduced in this mutant. Pyruvate concentrations were in fact the same in *cue1* and wild-type (Fig. 3b). The *cue1* mutant displays a reduced level of DXP compared to wild-type, but it also possesses a lower concentration of GAP (Fig. 3b), which affects the rate of the DXS reaction. When normalized to its corresponding GAP level, the DXP levels in *cue1* are unchanged compared to wild-type (Fig. 3b), indicating that PPT1 is unlikely to play a role in supplying pyruvate in adult leaf tissue. PPT1 is preferentially expressed in leaf vasculature and roots^33^, and our results support a likely role in providing PEP to the chloroplast in sink tissue.

**Fig 3.**
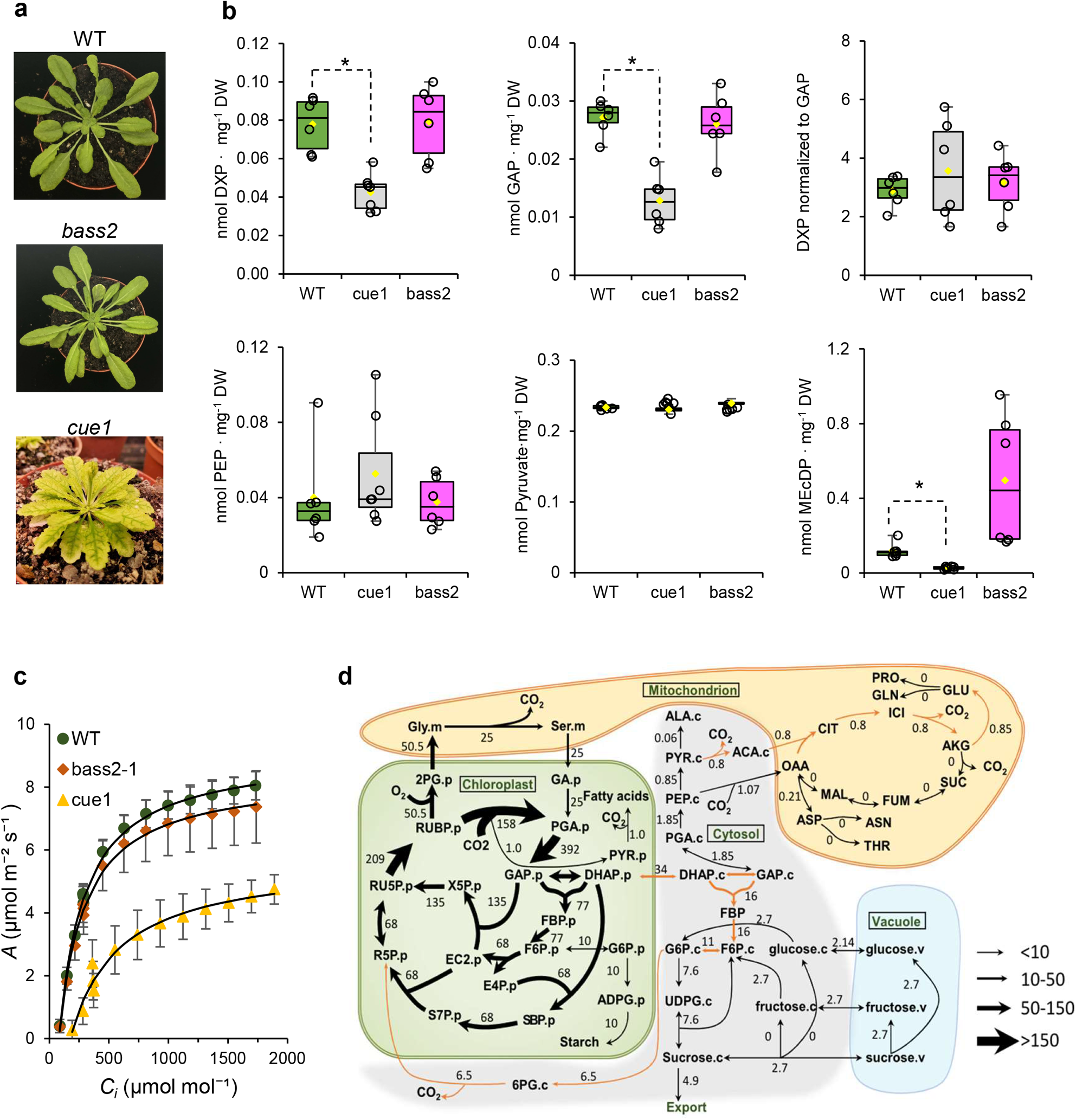
Role of transporters in supplying pyruvate to the chloroplast in Arabidopsis leaf tissue. **a**, Phenotypes of mutants in this study include the chlorophyll a/b-binding protein underexpressed 1 (*cue1*) mutant and the bile acid:sodium symporter family protein 2 (bass2) mutant. **b**, Absolute concentrations of selected metabolites in wild-type, *cue1* and *bass2* mutants. The data are visualized as box plots, with the central line indicating the median, upper and lower bounds represents quartile 3 (75th percentile) and quartile 1 (25th percentile), respectively, and the whiskers denotes the minimal and maximal values of the data points. Mean values (yellow diamond) are representative of n=6-7 individual plants. Asterisks indicate significant differences at * P < 0.01, based on Student’s two-tailed t-test. **c**, CO_2_ response curve showing CO_2_ assimilation (*A*) as a function of intercellular CO_2_ concentration (*C*_*i*_) in wild-type, *bass2* and *cue1* mutants. Curves were fitted based on the Farquhar-von Caemmerer-Berry (FvCB) model of photosynthesis (see supplementary figure 1). Data represent mean ± SD (n = 3). **d**. Flux model of central metabolism constructed with isotopically non-stationary metabolic flux analysis (see Supplementary Data Set 2 for additional details).

Direct import of pyruvate from the cytosol was also tested by measuring metabolite levels in the *bass2* mutant line, which lacks the pyruvate/Na^+^ symporter that supplies pyruvate in embryos^25^. The *bass2* mutant had wild-type levels of pyruvate, DXP, GAP, PEP, and MEcDP (Fig. 3b), suggesting that the MEP pathway does not depend on BASS2 for pyruvate import in autotrophic tissue. In addition to chloroplastic isoprenoids, pyruvate also supplies fatty acid biosynthesis and the branched chain amino acids leucine and valine in the chloroplast. AAAs were unchanged in the *bass2* mutant but reduced ∼40% in *cue1* rosette tissue (Supplementary Fig. 4), consistent with the role of PPT1 in supplying PEP for the shikimate pathway (it should be noted the *cue1* mutant requires AAA supplementation in the growth medium at 1 mM, and the observed levels may partly reflect low levels of AAAs retained following transfer to soil). However, we detected no changes in total fatty acid levels (Supplementary Fig. 5 and Supplementary Data Set 1) or branched chain amino acids (Supplementary Fig. 4) in either the *cue1* or *bass2* mutants, indicating that pyruvate availability in the chloroplast was not dependent on either PPT1 or BASS2.

A CO_2_ response (*A/Ci*) curve was performed to evaluate the physiological significance of these C_3_ translocators. Gas exchange measurements indicated that while the *A/C*_*i*_ response curve for *bass2* displayed slightly lower average *V*_*cmax*_ (maximal carboxylation rate) and *J*_*max*_ (electron transport rate) compared to wild-type (Supplementary Fig. 3), these differences were not statistically significant. However, in the *cue1* mutant, *V*_*cmax*_ was significantly lower than in wild-type (17.6 μmol·m^−2^·s^−2^ vs 28.5 μmol·m^−2^·s^−2^; n=3, p=0.018), consistent with its generally impaired growth aspect and pigment content (Fig. 3a and 3c).

Our results under low O_2_ conditions and in mutant lines identified rubisco β-elimination as the likely source of pyruvate in photosynthetic cells. We applied isotopically nonstationary metabolic flux analysis (INST-MFA) to further test this hypothesis. Using a previously reported photoautotrophic model of central metabolism^34^, we examined changes to goodness of fit when the model was adjusted to permit generation of pyruvate by rubisco. By permitting rubisco-mediated formation of pyruvate, our model predicted a rubisco-mediated pyruvate flux of 1.0 μmol.metabolite·g^−1^·FW·hr^−1^, with a corresponding flux into 3PGA of 158 μmol.metabolite·g^−1^·FW·hr^−1^ (Fig. 3d and Supplementary Data Set 2). This corresponds to a total rubisco pyruvate output of 0.63%, which closely aligns with our experimental measurement of 0.71% (Fig. 1f). Compared to a nearly identical metabolic model in which direct pyruvate formation by rubisco was not permitted, we observed a reduction in the sum of squared residues from 835 to 812 when pyruvate formation by rubisco was included (Supplementary Data Set 2). This modeling approach supports the experimental evidence implicating rubisco β-elimination as the principal source of pyruvate formation in the chloroplast.

Whole leaf extraction for metabolite analysis obscures cell specific differences in pyruvate trafficking. *PPT1* and *BASS2* are both expressed in developed leaves^35^, yet our metabolic and biochemical evidence clearly supports rubisco as the primary source of pyruvate on the whole leaf scale. These observations may be harmonized by considering cell specific expression of these translocators. *PPT1* is primarily expressed in vasculature tissue and roots^33^, while pyruvate sourcing through rubisco activity better explains its origin in mesophyll cells. The apparent contradiction of pyruvate labeling rates and pool size compared to PEP and other intermediates of the CBC has been described as the ‘Pyruvate Paradox’^4^. The significant downregulation of PGM and ENO in chloroplasts (Fig. 1f) precludes efficient pyruvate production for fatty acids and isoprenoids, and yet the *cue1* mutant does not show the expected symptoms of pyruvate deficiency in its fatty acid profile if PEP import on PPT1 were indeed the source of pyruvate (Supplementary Fig. 5). Our data instead support the *in vitro* conclusions published more than 30 years ago by Andrews and Kane^1^, namely, that the β-elimination of rubisco is the principal source of pyruvate in actively photosynthesizing leaves, a view supported by our updated in vitro biochemical characterization, labeling studies, mutant analysis, and metabolic modeling. This view simultaneously resolves the ‘Pyruvate Paradox’ and provides new possibilities for engineering terpenoid biosynthesis in the chloroplast given the direct connection between rubisco and the MEP pathway. For instance, the notion that rubisco directly governs nearly half the carbon entering the MEP pathway provides new impetus for understanding the regulation of rubisco by rubisco activase^36^ and other factors. Follow up studies aimed at single cell metabolome analyses and the impact of these insights on crop productivity and climate change mitigation strategies are currently underway.

## Online Methods

### Plant growth conditions

*Arabidopsis thaliana* (ecotype Columbia 0) plants were grown in 5 cm pots. Seeds were stratified in moist BX soil (Promix) for 3 days at 4 °C in the dark, then transferred to short day (SD) conditions that consisted of 8/16 hour light/dark cycles at 150 μmol photons·m^−2^·s^−1^, 21 °C, and 60-70% relative humidity (RH). Plants were fertilized once per week with MiracleGro™ (20/20/20) according to manufacturer’s instructions and otherwise watered as needed. Mutant lines *cue1*, and *bass2* were obtained from the Arabidopsis Biological Resource Center (https://abrc.osu.edu/). Plants were used in labeling and gas exchange experiments at 6-8 weeks of age prior to initiation of flowering. The *cue1* mutant was grown in magenta boxes for 1 month on sterile media containing¾ strength Murashige-Skoog medium supplemented with 1 mM Phe, Tyr, and Trp and then transferred to 5 cm pots as above.

### Characterization of mutant and transgenic plant lines

The genotype of mutant lines was confirmed by PCR using genomic DNA isolated according to Edwards et al^37^. A 5 ng aliquot of genomic DNA was used as template in a PCR consisting of Taq Master Mix (FroggaBio) diluted to 1X with forward and reverse primers (0.5 μM final concentration each), template, and water to a final volume of 20 μL. PCR genotyping was carried out for *bass2* mutants under the following thermocycler conditions: Initial denaturation at 96 °C for 2 min (1×), main amplification phase (96 °C × 20 s, 55 °C × 45 s, 72 °C × 60 s; 40×), and a final extension (72 °C × 5 min; 1×) Applied Biosystems Veriti thermocycler. The T-DNA mutant of BASS2 (At2G26900) was confirmed to be homozygous by performing PCR on genomic DNA with F (TTGGGTTTTCAAAACATCTGC) and R (TTACATGCGTCAGCTGAGTTG) primers for the wild-type (all primers listed 5’→3’). The mutant allele was screen with the same F primer and Lbl1.3 (ATTTTGCCGATTTCGGAAC) as R primer. The *cue1* genotype was confirmed by its phenotype and by PCR with F (ATGCAAAGCTCCGCCGTATTC) and R (TAAGCAGTCTTTGGCTTTGGCT) primers for the full length PPT1 gene (At5G33320) using cDNA as described below. PCR products were analyzed by gel electrophoresis on 1% agarose gels containing 5 μg·mL^−1^ ethidium bromide (Bioshop). For quantitative PCR analysis (QPCR), total RNA was extracted from fresh leaf frozen leaf tissue using the Maxwell RSC Plant RNA Kit (Promega) according to manufacturer’s instructions (https://www.promega.ca/). First-strand cDNA was synthesized from 2 μg of total RNA with a polydT primer using the SuperScriptIV Reverse Transcriptase Kit (Invitrogen) in a 20 μL reaction. cDNA products were diluted 1:10 in pure water prior to QPCR analysis in a BIO-RAD CFX96 Touch system. Relative quantification was accomplished using *APT1* and *RP2ls* reference genes as previously described^38^. QPCR F and R primers were GACTACAAAACCTCACCATCTTCAG and CAGACATGATGGCGCAGATT, respectively, for plastidic *ENO* (At1G74030) and ACCAATCAGTCGTTTCATTTGC and TGTAAAGCTCAGTGAAGAATCGATC for plastidic *PGM1* (At1G22170).

### Gas exchange and ^13^CO_2_ labeling

Measurement of gas exchange parameters in whole Arabidopsis plants was accomplished with a LI-COR 6800 photosynthesis system fitted with an interface for a custom chamber. Design and fabrication of the custom dynamic flow cuvette for combined gas exchange and administration of ^13^CO_2_ was described previously^39^. During the adaptation phase of the experiment, room air was adjusted to 400 μL·L^−1^ CO_2_ and 70% relative humidity by the LI-COR 6800 console and directed over a single Arabidopsis rosette enclosed in the cuvette at a flow rate of 1 L·min^−1^ (∼ 680 μmol·s^−1^) for gas exchange measurements (*A, C*_*i*_, and *g*_*s*_). For *A/C*_*i*_ measurements, CO_2_ concentrations were sequentially set at 400, 300, 200, 100, 50, 10, 400, 400, 600, 800, 1000, 1200, 1400, 1600, 1800, and 2000 μmol·mol^−1^. Data were recorded at each concentration after values of *A* and *C*_*i*_ were stable. Curves were fitted with the Farquhar, von Caemmerer and Berry (FvCB) model of photosytheic using the R ‘plantecophys’ package^40^. The estimated *V*_*cmax*_ (maximum rate of rubisco carboxylation) and *J*_*max*_ (maximum rate of electron transport) from individual *A/Ci* curve were subjected to a two-tailed t-test to assess significant differences between wild-type and mutants (p < 0.05). Light intensity and temperature were maintained at 150 μmol·m^−2^·s^−1^ and 21 °C, respectively. A 100% step change to the labeling atmosphere (20% O_2_, 400 μL·L^−1 13^CO_2_, ∼80% N_2_; Linde Canada) from a compressed air tank initiated the labeling. Compressed air flow of the labeling atmosphere was controlled by an Omega Engineering digital flow controller (FMA-2606A) at the same RH. The flow rate was temporarily raised to 3.5 L·min^−1^ until the original atmosphere had been >99% replaced, as judged by CO_2_ sensors. Atmospheric exchange was monitored individually for each labeling assay, and the labeling time was corrected based on the time to reach a 50:50 mixture of atmospheres (usually 2-3 s after opening the valve). Following 30 min of continuous gas exchange measurement during the adaptation phase, labeling assays were performed at intervals between 5 s and 1 h prior to flash freezing. For low O_2_ labeling experiments, plants were adapted in an atmosphere containing air with 1% O_2_ and 400 μL·L^−1^ CO_2_ (balance N_2_; Linde Canada) at 1 L·min^−1^ and 70% RH, then switched to an otherwise identical low O_2_ air source with 400 μL·L^−1 13^CO_2_. The humidity of compressed air was regulated by passing a variable fraction of the air flow through a custom humidifier tank. At the completion of the labeling phase, plants were quenched *in situ* with liquid nitrogen, ground to a fine powder, and lyophilized to dryness against a vacuum of 20 μbar for 24 h.

### Gas and liquid chromatography – mass spectrometry analysis

All steps in sample quenching, metabolite extraction, data acquisition, quality control, internal standardization, and batch correction were carried out according to the recommendations of Alseekh et al^41^. Analysis of ^13^C label incorporation into pyruvate and other small polar metabolites was carried out by gas chromatography – mass spectrometry (GC-MS) as previously described ^16^. The corresponding analysis of charged (phosphorylated) metabolites was performed by liquid chromatography tandem MS (LC-MS/MS) according to previous methods ^39^. The isotopolog masses monitored to calculate fractional labeling of each metabolite are listed in Supplementary Table 4. Analysis of fatty acid methyl esters (FAME) was performed as previously described^42^. For analysis of amino acids, ∼15 mg dry powdered leaf tissue was extracted in 500 μL methanol, centrifuged for 5 min at 10,000 g, and filtered through a 0.2 μm polyfluorotetraethylene filter prior to analysis by LC-MS/MS using settings in Supplementary Table 5.

### Data analysis and statistical methods

LC-MS/MS data analysis was conducted using Sciex OS (v2.0.0). GC-MS data analysis was performed with Agilent MassHunter Qualitative Analysis (v10.0). Metabolite peaks were initially validated by matching retention time and mass with those of authentic standards, then integration of extracted ion chromatograms (EIC) was performed to obtain retention time, mass spectra and peak area information. Total ^13^C labeled fraction of each metabolite was computed as previously described^28^. Time-course data were fitted to an exponential rise to maximum according to the equation A×(1-e^[-k × t]), where A is the labeling plateau, t is the labeling time, and *k* is the kinetic rate constant. Absolute concentrations of each metabolite in plant extracts were quantified using external calibration curves derived from authentic standards. Peak areas were normalized to internal standards and compared to the linear regression obtained from the calibration curves. Statistical significance was assessed using a two-tailed Student’s t test, with a significance cutoff set at P<0.05

### Biochemical characterization

Rubisco was purified from *Nicotiana tabacum* (tobacco) and activated in 50 mM EPPS buffer (EPPS 50 mM, NaHCO_3_ 15 mM, EDTA 0.2 mM, MgCl_2_ 20 mM, pH 8.0) for 5 min at 25 °C. The reaction is initiated by adding 0.5 mM RuBP into the reaction buffer at 25 °C and is allowed to proceed for 60 min, then terminated by rapidly freezing using liquid nitrogen. Samples were lyophilized to dryness and stored at −80 °C until subsequent LC-MS/MS and GC-MS analyses. Ribulose 1,5-bisphosphate and 3-phosphoglyceric acid were analyzed using ion-pair chromatography-tandem mass spectrometry (IPC-MS/MS) employing an ACQUITY UPLC pump system (Waters, Milford, MA, USA) coupled with a Waters XEVO TQ-S UPLC/MS/MS (Waters, Milford, MA, USA). Pyruvate underwent initial derivatization through methoximation followed by tert-butyldimethylsilylation. Subsequently, the derivatized pyruvate was analyzed using GC-MS, employing an Agilent 7890 GC system (Agilent, Santa Clara, CA, USA) coupled to an Agilent 5975C inert XL Mass Selective Detector (Agilent, Santa Clara, CA, USA) with an autosampler (CTC PAL; Agilent, Santa Clara, CA, USA). The IPC-MS/MS and GC-MS methods including MS parameters for metabolite measurements through multiple reaction monitoring (MRM) in IPC-MS/MS and selected ion monitoring (SIM) in GC-MS were as described previously^43,44^. IPC-MS/MS data was analyzed using MassLynx 4.0 (Agilent, Santa Clara, CA, USA). GC-MS data was analyzed with the Agilent GC/MSD ChemStation (Agilent, Santa Clara, CA, USA). Metabolite identification relied on retention time and mass-to-charge ratio (m/z), compared to authentic standards. Metabolites were quantified using calibration curves derived from external authentic standards through the QuanLynx software (Agilent, Santa Clara, CA, USA).

### Flux modeling

INST-MFA was conducted to estimate metabolic fluxes using the Isotopomer Network Compartmental Analysis software package (INCA2.1, Vanderbilt University)^45^, employing a previously established metabolic network model^43^. Goodness of fit, measured by sum of squared residuals (SSR), quantified the total difference between measured and simulated kinetics. 95% confidence intervals for fluxes were estimated by the parameter continuation method. Supplementary Dataset 2 provides a list of reactions and abbreviations, stoichiometry, atom transitions for each reaction, and 95% confidence intervals for fluxes. Computationally intensive tasks were parallelized through a SLURM job scheduler on a high-performance computing cluster at the Institute for Cyber-Enabled Research at Michigan State University (https://icer.msu.edu/).

## Acknowledgments

This work was supported by a grant from the Division of Chemical Sciences, Geosciences and Biosciences, Office of Basic Energy Sciences of the United States Department of Energy (Grant DE-FG02-91ER20021). TDS received partial salary support from Michigan AgBioResearch. Rubisco was isolated from *Nicotiana tabaccum* by Audrey Johnson. We also acknowledge help from the MSU Mass Spectrometry and Metabolomics Core Facility, MSU Institute for Cyber-Enabled Research for providing a high-performance computing cluster and services, and Dr. Jamey Young for making INCA accessible. This study was also funded by a Discovery grant from the Natural Sciences and Engineering and Research Council (NSERC) of Canada (RGPIN-2017-06400) and a John Evans Leadership Fund grant from the Canadian Foundation for Innovation (36131). The authors also acknowledge a generous NSERC CGSD graduate scholarship supporting MEB and thank UTM staff members Brenda Pitton of the Teaching Greenhouse, Peter Duggan of the Academic Machine Shop, and Stores Supervisor Matthew Malcom for their excellent technical and logistical support.

## Author contributions

SEE and AEF labeled plants and performed mass spectrometry analysis. SEE and MEB performed data analysis. SAF analyzed FAMEs and amino acids. YX measured pyruvate made by rubisco and carried out the mass flux analysis. TDS edited the manuscript. MAP wrote the manuscript and directed the research.

