## Supplementary Information for "Rubisco supplies pyruvate for the 2-*C*-methyl-D-erythritol-4-phosphate pathway"

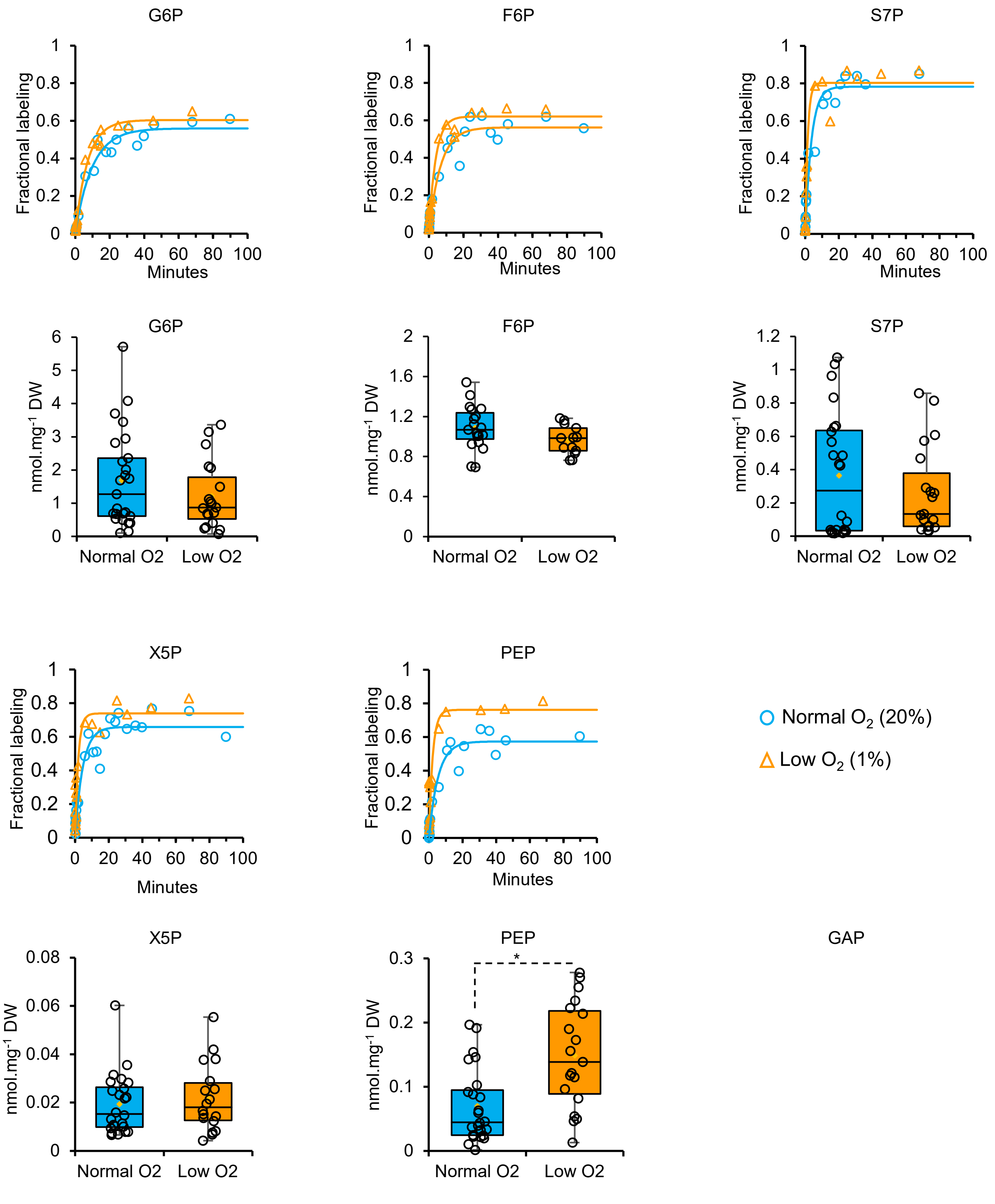


**Supplementary Fig 1. $^{13}$C labeling incorporation and absolute concentrations of glycolytic and Calvin-Benson-Bassham cycle intermediates under normal (21%) and low (1%) oxygen conditions**. For absolute quantification, the data are visualized as box plots, with the central line indicating the median, upper and lower bounds represents quartile 3 (75th percentile) and quartile 1 (25th percentile), respectively, and the whiskers denotes the minimal and maximal values of the data points. Mean values (yellow diamond) are representative of n=18-25 individual plants. Asterisks indicate significant differences at * $P < 0.01$, based on Student's two-tailed t-test. Glyceraldehyde-3-phosphate (GAP) labeling in time course assays represents labeling performed under normal oxygen conditions.

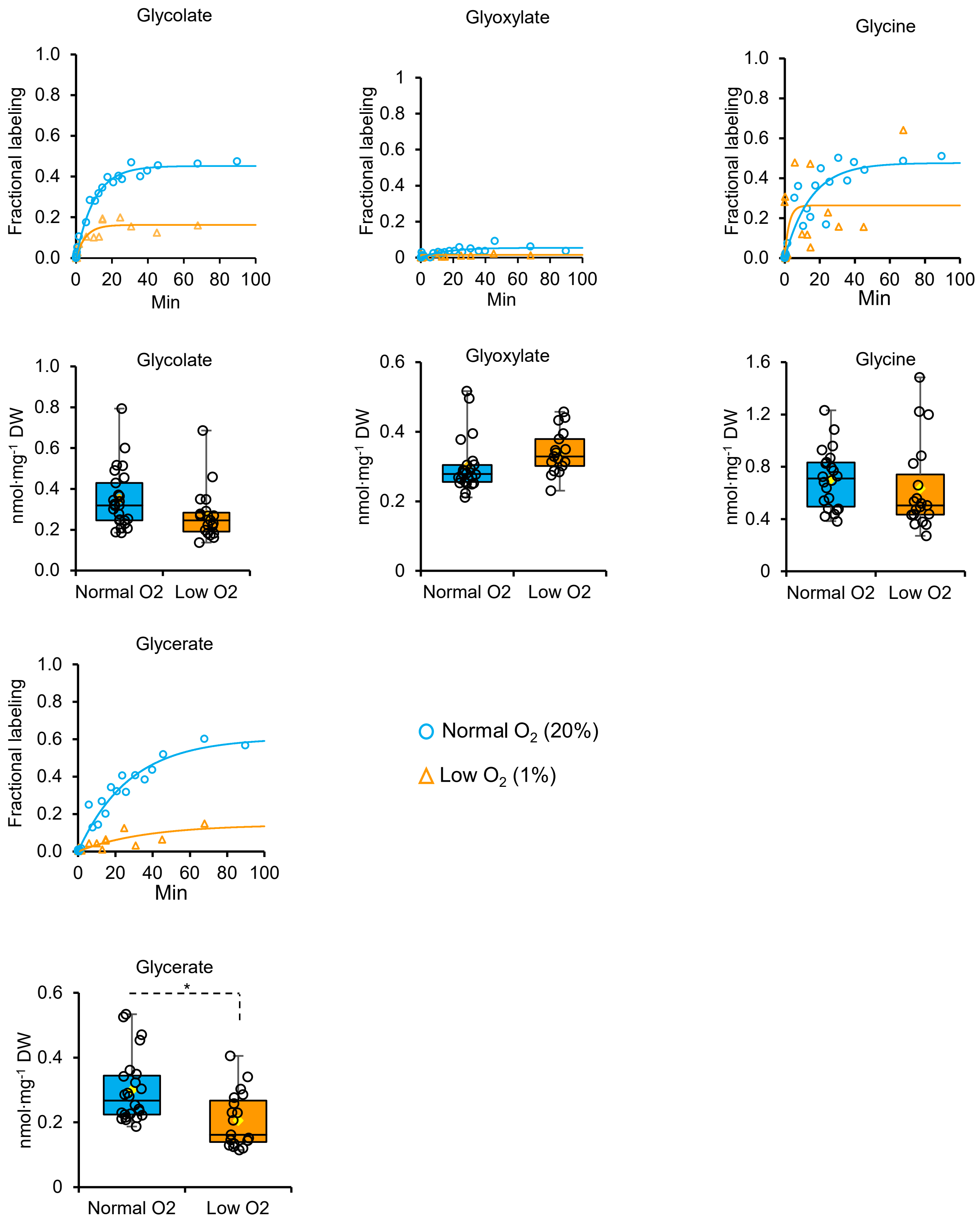


**Supplementary Fig 2. $^{13}C$ label incorporation and absolute concentrations of photorespiratory intermediates in Arabidopsis leaves under normal (21%) and low (1%) oxygen conditions.** Details on administration of 13C under physiological conditions, extraction of metabolites, and analysis by liquid chromatography – tandem mass spectrometry are given in the Methods section. For absolute quantification, the data are visualized as box plots, with the central line indicating the median, upper and lower bounds represents quartile 3 (75th percentile) and quartile 1 (25th percentile), respectively, and the whiskers denotes the minimal and maximal values of the data points. Mean values (yellow diamond) are representative of n=18-25 individual plants. Asterisks indicate significant differences at * $P < 0.01$, based on Student's two-tailed t-test.

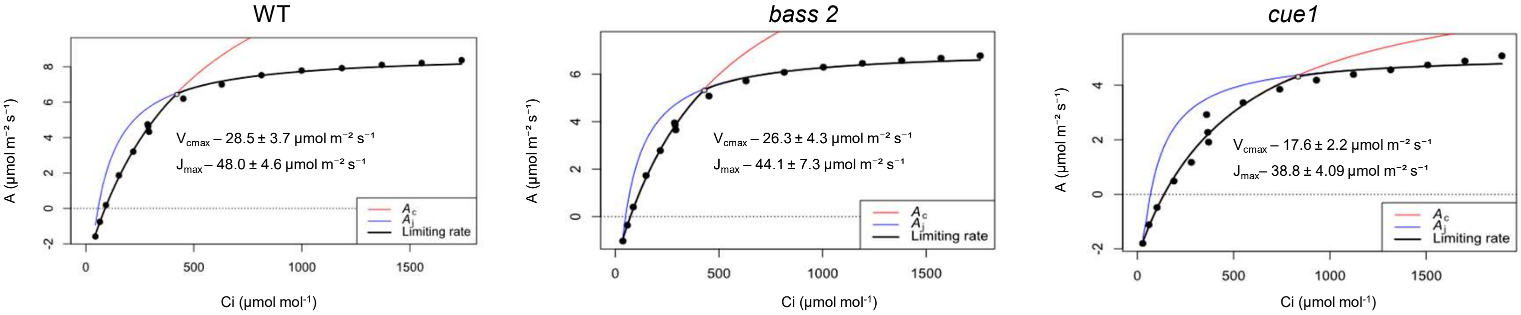


**Supplementary Fig 3. $CO_2$ assimilation (*A*) and intercellular $CO_2$ concentration (*$C_i$*) in Arabidopsis wild-type, *bass2* and *cue1* mutant plants.** *A/Ci* curves were fit based on the Farquhar-von Caemmerer-Berry (FvCB) model of photosynthesis using the plantecophys package on R[1]. Estimates of $V_{cmax}$ (maximum Rubisco carboxylation rate) and $J_{max}$ (the maximum rate of electron transport) are presented as mean ± SD from individual fit (n = 3 plants). $A_c$ represents photosynthesis limited by rubisco, while Aj indicates limitation due to RuBP-regeneration.

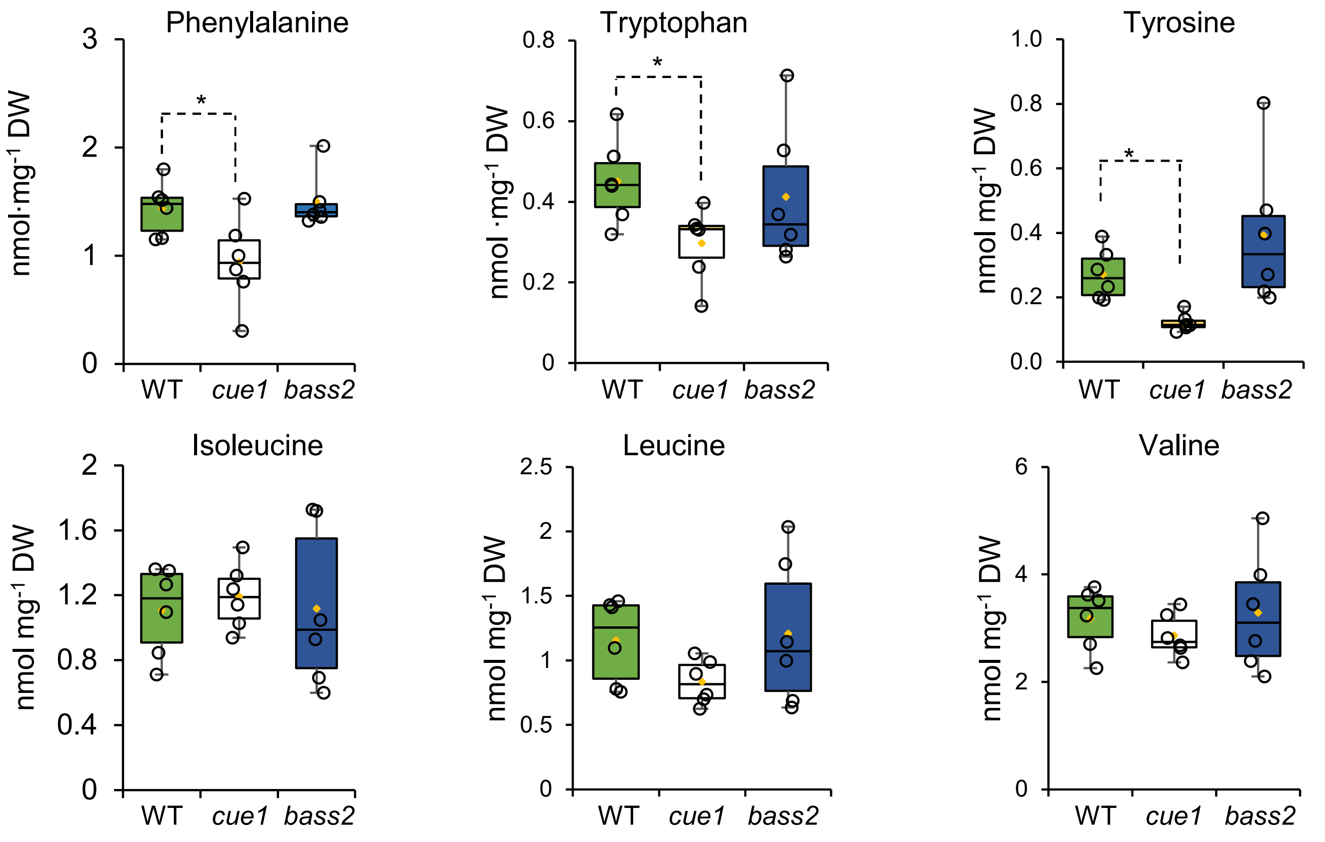


**Supplementary Fig 4. Roles of transporters in supplying pyruvate and phosphoenolpyruvate for amino acid biosynthesis in the chloroplast.** Concentrations of aromatic (phenylalanine, tryptophan, and tyrosine) and branched chain amino acids (isoleucine, leucine and valine) were analyzed by liquid chromatography – tandem mass spectrometry in leaf extracts from wild-type (WT), *cue1*, and *bass2* plants. Raw peak areas were normalized to the internal standard N-methylglucamine and compared to external standard calibration curves derived from authentic standards. The data are visualized as box plots, with the central line indicating the median, upper and lower bounds represents quartile 3 (75$^{th}$ percentile) and quartile 1 (25$^{th}$ percentile), respectively, and the whiskers denote the minimal and maximal values of the data points. Yellow diamonds indicate mean values (n=6-7 individual plants). Asterisk indicates significant differences at * $P < 0.05$, based on Student's two-tailed t-test.

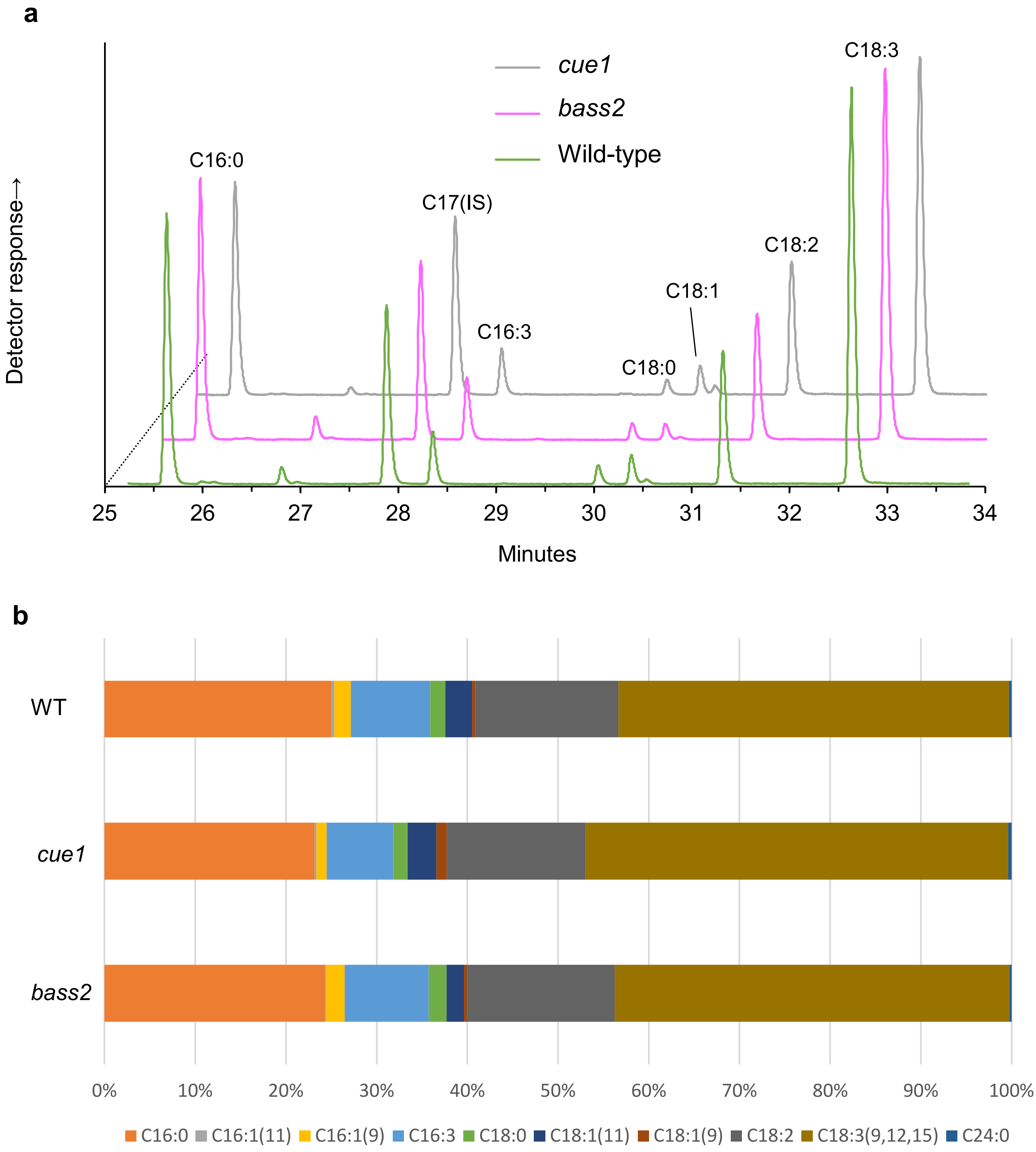


**Supplementary Fig 5. Fatty acid methyl ester (FAME) analysis in Arabidopsis wild-type (WT), *cue1*, and *bass2* plants. a**, Representative chromatogram showing elution profile of FAMEs by gas chromatography – mass spectrometry. **b**, Stacked plot showing relative distribution of fatty acids in these same genotypes. These results show the average values (n = 4) for FAME analysis using triheptadecanoin (C17) as internal standard (see Methods for additional details). There were no statistical differences in total fatty acid content between groups, but content of minor components (C16:1(9) and C18:1(9)) making up <1% of the total FAME peak area were lower in cue1 (see Extended Data Set 1 for details).

**Supplementary Table 1. Pyruvate production by rubisco.** Assays were incubated with or without rubisco for 60 min at 25°C. Samples were processed for LC-MS/MS as described in the methods. Recovery of RuBP (+PGA and pyruvate assuming these are spontaneously produced from RuBP) without rubisco was 88.0% while recovery with rubisco calculated as (RuBP +(PGA + pyruvate)/2)/63.5 was 92.8%. The difference was not statistically significant at the 5% level by t-test in Excel. Pyruvate and PGA in the minus rubisco treatment may have been contamination or spontaneous conversion. Subtracting these amounts from the PGA and pyruvate in the + rubisco samples and taking (PGA + pyruvate)/ 2 to be total carboxylations, there were an average of 0.71 pyruvates per 100 carboxylations. The small number of samples gives a large 95% confidence interval of 0.57 to 0.85% (calculated in Excel). Since there are three carbons per pyruvate, pyruvate can account for 2% of the carbon leaving the Calvin-Benson cycle. RuBP = ribulose 1,5-bisphosphate, PGA = 3-phosphoglyceric acid

| Replicate | RuBP added | RuBP recovered | PGA | Pyruvate | Pyruvate/ carboxylation |
|---|---|---|---|---|---|
| + Rubisco | (μM) | (μM) | (μM) | (μM) | % |
| 1 | 63.5 | 0.16 | 112 | 0.35 | 0.57 |
| 2 | 63.5 | 0.12 | 117 | 0.55 | 0.90 |
| 3 | 63.5 | 0.12 | 124 | 0.42 | 0.63 |
| 4 | 63.5 | 0.12 | 115 | 0.44 | 0.73 |
| **Average** | **63.5** | **0.13** | **117** | **0.44** | **0.71** |
| - Rubisco | | | | | |
| 1 | 63.5 | 53.7 | 0.65 | 0.026 | |
| 2 | 63.5 | 55.8 | 0.58 | 0.021 | |
| 3 | 63.5 | 57.8 | 0.57 | 0.029 | |
| 4 | 63.5 | 55.0 | 0.67 | 0.025 | |
| **Average** | **63.5** | **55.6** | **0.62** | **0.025** | |

**Supplementary Table 2.** Kinetics parameters of $^{13}C$ incorporation into organic acids of photorespiration and primary metabolism. Data was fitted to an exponential rise to maximum equation $[A \times (1 - e^{[-k \times t]})]$, where A represents the labeling plateau, $t$ is the labeling time, and $k$ is the kinetic rate constant.

| **Low O2** | Glyoxylate | Pyruvate | Glycolate | Glycine | Glycerate |
|---|---|---|---|---|---|
| *A* | 0.016 | 0.318 | 0.162 | 0.264 | 0.143 |
| *k* | 9.926 | 0.064 | 0.186 | 0.405 | 0.028 |
| $Chi^2$ | 0.008 | 0.005 | 0.011 | 0.524 | 0.011 |

| **Normal O2** | Glyoxylate | Pyruvate | Glycolate | Glycine | Glycerate |
|---|---|---|---|---|---|
| *A* | 0.054 | 0.236 | 0.452 | 0.475 | 0.606 |
| *k* | 0.063 | 0.049 | 0.098 | 0.068 | 0.036 |
| $Chi^2$ | 0.004 | 0.037 | 0.010 | 0.133 | 0.042 |

**Supplementary Table 3**. Kinetics parameters of $^{13}C$ incorporation into MEP pathway, glycolysis, and CBC intermediates in Arabidopsis wild-type labeled under normal (21%) and low (1%) oxygen. Isotopolog analysis was conducted using LC-MS/MS. Data was fitted to an exponential rise to maximum equation $[A \times (1 - e^{[-k \times t]})]$, where A represents the labeling plateau, $t$ is the labeling time, and $k$ is the kinetic rate constant. Parentheses indicate downstream intermediates whose rate constants can only be estimated with this method. 3PGA, 3-phosphoglycerate; Triose-P, triose phosphate (glyceraldehyde-3-phosphate and dihydroxyacetone phosphate); G6P, glucose 6-phosphate; S7P, sedoheptulose 7-phosphate; DXP, deoxyxylulose 5-phosphate; MEcDP, methylerythritol cyclodiphosphate; IDP/DMADP, isopentenyl diphosphate and dimethylallyl diphosphate (cannot be resolved chromatographically by this method).

| Low O2 | Triose-P | 3PGA | PEP | Xu5P | G6P | F6P | S7P | DXP | MEcDP | IDP/DMADP |
|---|---|---|---|---|---|---|---|---|---|---|
| *A* | 0.761 | 0.887 | 0.762 | 0.739 | 0.596 | 0.621 | 0.783 | 0.486 | 0.666 | 0.743 |
| *k* | 0.523 | 0.605 | 0.444 | 0.530 | 0.140 | 0.256 | 0.275 | 0.264 | (0.106) | (0.097) |
| $Chi^2$ | 0.080 | 0.065 | 0.269 | 0.168 | 0.008 | 0.025 | 0.097 | 0.097 | 0.025 | 0.055 |

| Normal O2 | | | | | | | | | | |
|---|---|---|---|---|---|---|---|---|---|---|
| *A* | 0.690 | 0.799 | 0.572 | 0.659 | 0.550 | 0.563 | 0.804 | 0.494 | 0.558 | 0.674 |
| *k* | 0.218 | 0.418 | 0.183 | 0.221 | 0.101 | 0.148 | 0.695 | 0.121 | (0.133) | (0.087) |
| $Chi^2$ | 0.208 | 0.049 | 0.067 | 0.152 | 0.029 | 0.059 | 0.059 | 0.099 | 0.0271 | 0.098 |

**Supplementary Table 4**. Mass isotopologs used in the calculation of fractional labeling of each metabolite pool. Phosphorylated metabolites were analyzed by liquid chromatography – tandem mass spectrometry (LCMS/MS) using a Sciex 4500 Qtrap operating in multiple reaction monitoring (MRM) in negative mode. Q1, *m/z* of precursor ion; Q3, *m/z* of product ion. Gas chromatography – mass spectrometry (GCMS) analysis was performed using ammonia chemical ionization to minimize fragmentation (see Methods for details).

**LCMS/MS parameters**

| Metabolite | C atoms | Precursor range | Q1[Q3] (*m/z*) mass transitions |
|---|---|---|---|
| 3PGA | 3 | $[M-H]^-$ - $[M-H+3]^-$ | 185[79], 186[79], 187[79], 188[79] |
| Triose-P | 3 | $[M-H]^-$ - $[M-H+3]^-$ | 169[97], 170[97], 171[97], 172[97] |
| PEP | 3 | $[M-H]^-$ - $[M-H+3]^-$ | 167[79], 168[79], 169[79], 170[79] |
| X5P | 5 | $[M-H]^-$ - $[M-H+5]^-$ | 229[97], 230[97], 231[97], 232[97], 233[97], 234[97] |
| G6P | 6 | $[M-H]^-$ - $[M-H+6]^-$ | 259[97], 260[97], 261[97], 262[97], 263[97], 2644[97], 265[97] |
| F6P | 6 | $[M-H]^-$ - $[M-H+6]^-$ | 289[97], 290[97], 291[97], 292[97], 293[97], 294[97], 295[97], 296[97] |
| S7P | 7 | $[M-H]^-$ - $[M-H+7]^-$ | 229[97], 230[97], 231[97], 232[97], 233[97], 234[97] |
| DXP | 5 | $[M-H]^-$ - $[M-H+5]^-$ | 213[79], 214[79], 215[79], 216[79], 217[79], 218[79] |
| MEcDP | 5 | $[M-H]^-$ - $[M-H+5]^-$ | 277[79], 278[79], 279[79], 280[79], 281[79], 282[79] |
| IDP/DMADP | 5 | $[M-H]^-$ - $[M-H+5]^-$ | 245[79], 246[79], 247[79], 248[79], 249[79], 250[79] |

**GCMS parameters**

| Metabolite | C atoms | Adduct | Isotopolog series | Derivatization[a] |
|---|---|---|---|---|
| Pyruvate | 3 | $[M + NH_4]^+$ | 207, 208, 209, 210 | MeOX/TMS |
| Glycerate | 3 | $[M + NH_4]^+$ | 340, 341, 342, 343 | 3TMS |
| Glycolate | 2 | $[M + NH_4]^+$ | 238, 239, 240 | 2TMS |
| Glyoxylate | 2 | $[M + NH_4]^+$ | 193, 194, 195 | MeOX/TMS |
| Glycine | 2 | $[M + H]^+$ | 292, 293, 294 | 3TMS |

[a] MeOX, methyloxime derivative generated with methoxylamine. TMS, trimethyl silyl derivative formed with *N*-methyl-*N*-(trimethylsilyl)trifluoroacetamide.

**Supplementary Table 5**. Parameters for quantification of central metabolites by liquid chromatography – tandem mass spectrometry

| MS parameters used in LCMS/MS multiple reaction monitoring (MRM) | | | | | | | |
|---|---|---|---|---|---|---|---|
| **Metabolite** | **Precusor ion (Q1)** | **Product ion (Q3)** | **ESI polarity** | **DP** | **EP** | **CE** | **CXP** |
| | **(*m/z*)** | **(*m/z*)** | | **(v)** | **(v)** | **(v)** | **(v)** |
| PGA | 185 | 79 | (-) | -23 | -10 | -41 | -10 |
| Triose-P | 169 | 97 | (-) | -35 | -10 | -20 | -20 |
| PEP | 167 | 79 | (-) | -5 | -10 | -14 | -5 |
| Xu5P | 229 | 97 | (-) | -5 | -10 | -18 | -6 |
| G6P | 259 | 97 | (-) | -17 | -8 | -19 | -9 |
| F6P | 259 | 97 | (-) | -31 | -8 | -58 | -11 |
| S7P | 289 | 97 | (-) | -38 | -5 | -22 | -7 |
| DGP[a] | 243 | 79 | (-) | -35 | -10 | -62 | -7 |
| DXP | 213 | 79 | (-) | -30 | -10 | -40 | -8 |
| MEcDP | 277 | 79 | (-) | -30 | -10 | -65 | -11 |
| IDP/DMADP | 245 | 79 | (-) | -45 | -6 | -24 | -6 |
| Tryptophan | 205 | 188 | (+) | 41 | 9 | 19 | 9 |
| Phenylalanine | 166 | 103 | (+) | 10 | 8 | 35 | 10 |
| Tyrosine | 182 | 136 | (+) | 45 | 10 | 19 | 10 |
| Isoleucine | 132 | 69 | (+) | 70 | 10 | 25 | 12 |
| Leucine | 132 | 43 | (+) | 70 | 10 | 35 | 12 |
| Valine | 118 | 72 | (+) | 40 | 10 | 5 | 12 |
| N-methylglucamine[b] | 196 | 178 | (+) | 60 | 4 | 21 | 12 |

[a] DGP was used as internal standard for phosphorylated metabolites.

[b] N-methylglucamine was used as internal standard for amino acid quantification

ESI, electrospray ionization; DP, declustering potential; EP, entrance potential; CE, collision energy; CXP, collision cell exit potential; V, volts. Compound abbreviations: PGA, 3-phosphoglycerate; Triose-P, triose phosphate (glyceraldehyde-3-phosphate and dihydroxyacetone phosphate); PEP, phosphenolpyruvate; Xu5P, xylulose-5-phosphate; G6P, glucose-6-phosphate; F6P, fructose-6-phosphate; S7P, sedoheptulose-7-phosphate; DXP. 1-deoxyxylulose-5-phosphate; MEcDP, 2-C-methylerythritol-2,4-cyclodiphosphate, IDP, isopentenyl diphosphate; DMADP, dimethylallyl diphosphate.
